# CP_dock_: The Complementarity Plot for Docking of Proteins: Implementing Multi-dielectric Continuum Electrostatics

**DOI:** 10.1101/185686

**Authors:** Sankar Basu

## Abstract

The Complementarity plot (CP) is an established validation tool for protein structures, applicable to both, globular proteins (folding) as well as protein-protein complexes (binding). It computes the shape and electrostatic complementarities (S_m_, E_m_) for amino acid side-chains buried within the protein interior or interface and plots them in a two-dimensional plot having knowledge-based probabilistic quality estimates for the residues as well as for the whole structure. The current report essentially presents an upgraded version of the plot with the implementation of the advanced multi-dielectric functionality (as in Delphi version 6.2 or higher) in the computation of electrostatic complementarity to make the validation tool physico-chemically more realistic. The two methods (single‐ and multi-dielectric) agrees decently in their resultant E_m_ values and hence, provisions for both methods have been kept in the software suite. So to speak, the global electrostatic balance within a well-folded protein and / or a well-packed interface seems only marginally perturbed by the choice of different internal dielectric values. However, both from theoretical as well as practical grounds, the more advanced multi-dielectric version of the plot is certainly recommended for potentially producing more reliable results. The report also presents a new methodology and a variant plot, namely, CP_dock_, based on the same principles of complementarity, specifically designed to be used in the docking of proteins. The efficacy of the method to discriminate between good and bad docked protein complexes have been tested on a recent state-of-the-art docking benchmark. The results unambiguously indicate that CP_dock_ can indeed be effective in the initial screening phase of a docking scoring pipeline before going into more sophisticated and computationally expensive scoring functions. CP_dock_ has been made available at https://github.com/nemo8130/CPdock

## Introduction

The Complementarity plot (CP) is an existing graphical structure validation tool in the field of protein science originally proposed for globular proteins [1–3] and later extended for experimentally solved protein co-complexes [4]. So far, this has been constructed and applied to analyze complementarity in a residue-wise manner in proteins. CP has a wide array of applications ranging from serving as a component in protein crystallography, homology modeling, protein design and dynamics [1,2,5]. It computes the shape and electrostatic complementarities for amino acid side-chains deeply or partially buried within the protein interior or interface and plots these ordered pair values in a two-dimensional plot (E_m_: ordinate, S_m_: abscissa) according to the burial of the residues [1]. In effect, the term CP stands out to be a misnomer, while in reality, there exist three plots (CP1, CP2, CP3) corresponding to three different burial bins (see **Methods**) [1]. The design of CP is analogous to the famous Ramachandran Plot (RP) [6] though it is distinctly different in its physico-chemical nuances and accordingly in its particular uses in the validation pipeline. For example, structures with incorrectly assigned side-chain rotamers (frequently found in low-resolution atomic models) may indeed give rise to a sub-optimal distribution of points in their CPs, in spite of well satisfying the Ramachandran validation filters (or scores based on the RP – as in Procheck [7], Molprobity [8]) for having an optimal set of Φ, Ψ_s_ [2]. For this very reason, the combined application of the two complementarity measures (S_m_, E_m_) has also found its wide use and superiority in performance to older methods in threading exercises [1] wherein the problem was to identify the correct native structure situated amid a set of decoys, with their main-chains being kept identical to that of the native (offering an identical set of Φ, Ψs) while allowing variations only in their side-chain coordinates. Also, in the related problem of protein fold recognition, the methodology could prove its efficacy to identify the correct fold for a pair of sequences having low sequence similarity although belonging to the same fold [1].

The accurate determination of the local internal dielectric constant within proteins (as a function of the degree of solvent exposure of amino acid residues) has been a well posed problem in the field for decades. For electrostatic continuum models, there has been much debate among different values proposed for the parameter [9]. The subject was made further complicated by paradoxical observations like the presence of ionic groups in the hydrophobic protein interior without having to adapt to any specialized structures to get stabilized, demanding a considerably higher effect experienced locally at these sites than can be accounted for by the low dielectric constant traditionally assigned to represent the hydrophobic protein interior in continuum models [10]. Primarily contributed by this ‘dielectric problem’, electrostatic complementarity of individual amino acid residues (E_m_) within a folded protein chain remained undetermined, until the problem was finally adequately addressed by the same study [1] that put forward the Complementarity Plot (CP). Briefly, the study had shown that electrostatic complementarity (being a correlation function between two troughs of potential values) is independent of any single dielectric constant assigned to the protein interior, treated as the internal dielectric of the continuum [1]. Even then, the methodology had room to implement more sophistication by means of actually quantifying the local internal dielectric *in situ* prior to calculating the electrostatic potential. This central concern of protein electrostatics has precisely been addressed recently in an advanced version of Delphi [11], the leading Poisson-Boltzmann solver in the field. More specifically, by means of their Gaussian-based approach in the multi-dielectric treatment of the protein interior [12], Delphi can now deliver the dielectric constant distribution throughout a protein. As a result, it presented a realistic opportunity to incorporate this functionality in CP and recompute E_m_ as a function of variable local internal dielectric, compare the results with the earlier method, and keep both options in the user interface of the software and this is the central theme of the current report. The study shows strong agreement in general in the E_m_ values for amino acid residues within proteins, computed by both single and multi-dielectric methods. It is to be noted that large disagreement between the E_m_ values is rare and highly contextual, and, in such cases the more advanced multi-dielectric method is surely recommended. Subsequently, both softwares for CP (**Interior:** https://github.com/nemo8130/SARAMA-updated [3], **Interface:** https://github.com/nemo8130/SARAMAint-updated [4]) have been added with the provisions to implement both single and multi-dielectric methods as separate standalone packages.

In addition, here we also present a varient of the Complementarity Plot (namely, CP_dock_) to be effectively used at the initial screening phase of a protein-protein docking scoring pipeline. CP_dock_ is based on single values of shape (Sc) and electrostatic complementarity (EC) raised at the protein-protein interface, as was in the original formulations of the two complementarity measures [13,14].

## Methods

### Databases

For comparing the single‐ and multi-dielectric methods in terms of their effect in CP, the S_m_, E_m_ values for individual buried / partially buried residues were calculated from a previously standardized database (**DB2**) of globular proteins: https://github.com/nemo8130/DB2 which was used successfully in several earlier studies [1,2,15].

For the construction of CP_dock_, the Sc, EC values raised at protein-protein interfaces were calculated from a previously standardized database (DB3: https://github.com/nemo8130/DB3), used successfully in several other studies [4,15,16].

### The Complementarity Plot

Though the construction and implementation of CP has previously been described in complete detail [1–4], it is perhaps good to recapitulate some of its fundamental and essential features to make the current report self-contained. CP requires the shape (S_m_) and electrostatic (E_m_) complementarity to be computed for amino acid residues completely or partially buried within a folded polypeptide chain.

To start with, solvent accessible surface area (ASA) was calculated for each protein atom by NACCESS [17] which were then summed up for all atoms pertaining to the same residue. As elaborated in a series of earlier studies [1,2,18,19], the burial (*bur*) of solvent exposure for any residue (X) embedded in a polypeptide chain was defined by the solvent accessible surface area (ASA) of X located in the protein divided by the ASA of the same amino acid in a Gly-X-Gly peptide fragment in its fully extended conformation. Residues were distributed in three different burial bins based on their *bur* value mapping to three different CPs: CP1: 0.00 ≤ *bur* ≤ 0.05; CP2: 0.05 < *bur* ≤ 0.15; CP3: 0.15 < *bur* ≤ 0.30. The completely exposed residues with *bur >* 0.3 [18] were not considered in any of the CPs for lacking any notable packing constraints in their local neighborhood. For protein-protein interfaces, the atoms getting buried upon complexation, characterized by a net non-zero change in their corresponding ASA values in unbound and bound forms (i.e., |∆ASA| > 0) constituted the interface [4]. The corresponding amino acid residues were then distributed according to their burial and used to construct the corresponding CPs.

The van der Waals surface was then calculated for the entire polypeptide chain, sampled at 10 dots / Å^2^ [18] and the dot surface points sourced from each amino acid residue in the chain, identified. For shape complementarity (S_m_), only side-chain dot surface points corresponding to the buried residues (targets) were considered and their nearest neighboring surface points identified from the rest of the polypeptide chain (within a distance of 3.5 Ǻ). Surface points are essentially area elements and thus represented by their positions (x, y, z) and the direction cosines (dl, dm, dn) of their normals. Following Lawrence and Colman [13], the following expression was then calculated:

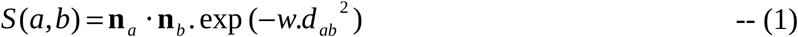

where **n***_a_* and **n***_b_* are two unit normal vectors, corresponding to the dot surface point *a* (located on the side-chain surface of the target residue) and *b* (the dot point nearest to *a*, within 3.5 Å) respectively, with *d_ab_* the distance between them and *w*, a scaling constant set to 0.5. S_m_ was defined as the median of the distribution {S(a,b)} calculated over all the dot surface points of the side-chain target residue.

Subsequent to identifying the nearest neighbors, the side-chain dot surface points of the specified residue (target) was partitioned into two sets by virtue of their neighbors coming from either side-chain or main-chain atoms, and calculate S_m_ separately for each set. Thus, every target residue (side-chain) has three measures of S_m_ based on the choice of its nearest neighbors (surface points), whether obtained from side-chain 
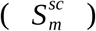
 or main-chain atoms 
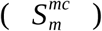
 alone, or all atoms (
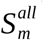
). Since, glycines lack any non-hydrogen side-chain atom, they were excluded as targets from all calculations.

For electrostatic complementarity (E_m_), the electrostatic potential of the molecular surface was estimated using the finite difference Poisson-Boltzmann method as in DelPhi [11] with the implementation of its advanced multi-dielectric features [12]. The potential on the side-chain surface points of a buried residue was then computed twice, firstly, due to all (charged) atoms of the target residue and secondly, due to all (charged) atoms from the rest of the polypeptide chain (excluding the target). Thus, each dot surface point was now tagged with two values of electrostatic potential. Following McCoy et al., [14], negative of the Pearson’s correlation between these two troughs of potential values over the dot surface points of the target residue was defined as E_m_.

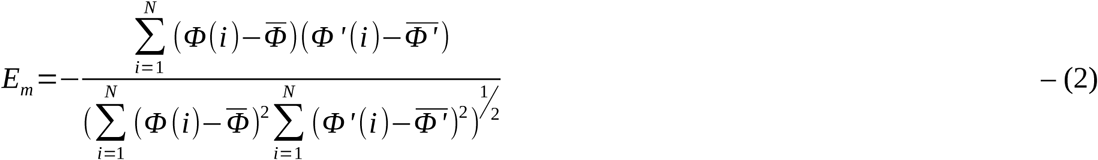

where, for a given residue consisting of a total of N dot surface points, Φ(i) is the potential on its ith point realized due to its own atoms and Φ′(i), due to the rest of the protein atoms, *Φ̅* and *Φ′̅* are the mean potentials of Φ(i), i = 1…N and Φ'(i), i = 1…N respectively.

Likewise to S_m_, subsequent to calculating the electrostatic potentials, the values corresponding to N dot surface points were divided into two distinct sets based on whether the dot point was obtained from main-chain or side-chain atoms of the target residue, and accordingly E_m_ was calculated separately for each set. Thus for a given residue, E_m_ was estimated for the entire residue (
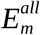
, as described above), the side-chain dot surface points 
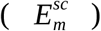
, and the main-chain dot surface points (
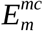
).

As discussed in the original reports [1,2], the plot of 
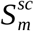
 on the X-axis and 
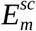
 on the Y-axis (spanning -1 to 1 in both axes) constitutes the ‘Complementarity Plot’ (CP), which is actually divided into three plots based on the burial ranges: 0.00 ≤ *Bur* ≤ 0.05 (CP1), 0.05 < *Bur* ≤ 0.15 (CP2) and 0.15 < *Bur* ≤ 0.30 (CP3). First, all the buried residues from the database, **DB2** were plotted in the CPs, which had been divided into square-grids (of width 0.05 × 0.05), and the center of every square grid was assigned an initial probability (P_grid_) equal to the number of points in the grid divided by the total number of points in the plot. The probability of a residue to occupy a specific position in the plot was then estimated by bilinear interpolation from the probability values of its four nearest neighboring voxels. Each CP was contoured based on the initial probability values (P_grid_ ≥ 0.005 for the first contour level and P_grid_ ≥ 0.002 for the second) thus dividing the plot into three distinct regions. Likewise to the ‘allowed', ‘partially allowed’ and ‘disallowed’ regions of the Ramachandran Plot [6], the region within the first contour was termed ‘probable’, between the first and second contour, ‘less probable’ and outside the second contour, ‘improbable’. An identical methodology was followed to probe protein-protein interfaces in a residue-wise manner [4].

It is expected from the distribution of points corresponding to residues coming from a well folded globular protein or a well-packed interface that most points should map to ‘probable’ and ‘less probable’ regions of the plot(s), while only a low fraction falling into the ‘improbable’ region, corresponding to local packing defects and/or sub-optimal electrostatic complementarity. This has been well demonstrated on thousands of high-resolution native X-ray structures of both folded globular proteins as well as packed interfaces [1–4].

For CP_dock_, the equivalent single-valued complementarity measures for protein-protein interfaces, namely, Sc and EC were directly adapted from their original formulations [13,14]. Sc was calculated precisely according to a previous report [16] while for EC, the only alteration in the previous protocol [16] was the permanent adaptation of the multi-dielectric continumm model [12].

### Software Implementation

The multi-dielectric versions of CP (**Interior:** https://github.com/nemo8130/SARAMA-updated/tree/master/SARAMA-multidielectric-delphi, **Interface:** https://github.com/nemo8130/SARAMAint-updated/tree/master/SARAMAint-multidielectric-delphi) have appropriately been added with the functionality to implement the delphi multi-dielectric model [12]. For user-convenience as well as for future development, the single and multi-dielectric methods have been kept as separate standalone packages; and, as a consequence, there is no alteration required at the user interface. The user may simply download the desired version (single‐ or multi-dielectric) and run an identical set of commands (as detailed in the software-documentation) to execute either of them. The CP packages (SARAMA, SARAMAint) can also be found enlisted in the delphi tools page (http://compbio.clemson.edu/delphi_tools) redirecting to their source site (http://www.saha.ac.in/biop/www/sarama.html) having the same currently updated versions and also hyperlinks to the aforementioned github pages. For all future downloads and updates, we recommend the user to refer to the github pages.

## Results and Discussion

### Statistical Comparison of the single‐ and multi-dielectric methods

The overall distribution of points in the CPs doesn’t really change by adapting to the multi-dielectric Gaussian model [12]. This was evident from the visual comparison of both distributions side-by-side across all three burial bins (**Fig.1, Supplementary Fig.S1, Fig.S2**). The agreement in the two methods was also quantified by the overlap in grid-probabilities (P_grid_) [1] between the corresponding pair of plots. P_grid_ values were found to be 0.94, 0.93, 0.92 calculated over 22461, 10997 and 13767 points for CP1, CP2 and CP3 for the interior (calculated on **DB2**) while almost identical overlap-values obtained also for the interface CPs: 0.94, 0.94, 0.93 for 27479, 18376, 17868 points (calculated on **DB3**). In accordance, the corresponding pair of E_m_ values (calculated by the two methods) also had descent agreement between themselves as revealed from their root mean square deviations (rmsd’s) and pair-wise Correlation (Pearson’s). The numbers were found to be rmsd: 0.12, Pearson’s: 0.90 (for 27479 residues in CP1), rmsd: 0.13, Pearson’s: 0.88 (for 18376 residues in CP2) and (rmsd: 0.15, Pearson’s: 0.86) for 17868 residues in CP3, calculated from the larger (**DB3**) of the two databases. Note, that the corresponding agreement between the equivalent single-valued term (EC) raised at protein-protein interfaces has already been reported (rmsd: 0.15, Pearson’s Correlation: 0.94) in a previous study [16].

**Fig.1.**
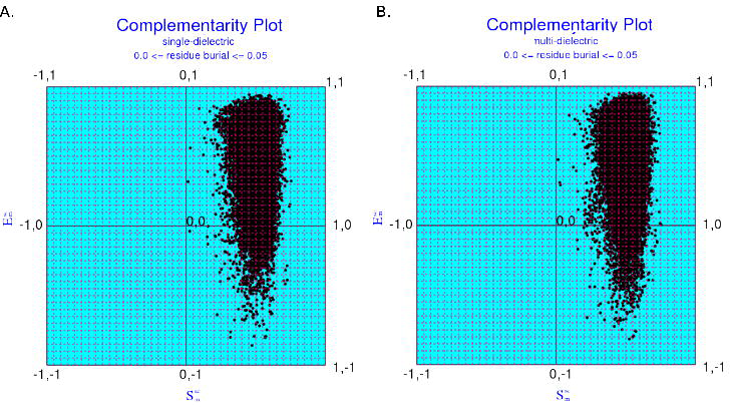
Distribution of points in CP1 (corresponding to the first burial bin: 0 ≤ *bur* ≤ 0.05) where E_m_ calculated by (A) single‐ and (B) multi-dielectric methods. S_m_, E_m_ calculated for amino acid residues from the database DB2. The distributions show the agreement between the two methods.

The contours to delineate the ‘probable', ‘less probable’ and ‘improbable’ regions [1] of the plots based on the new (multi-dielectric) distributions were also redrawn for all three CPs (**Fig.2**, **Supplementary Fig.S3, Fig.S4**) following an identical method described in the original formulation of CP [1] and compared with the earlier contours [1,2]. The contours were largely similar with only marginal increase in length along the Y-axis (E_m_) which is well within statistical limits of error. These ever-so-tiny differences could potentially also occur even if an identical methodology is implemented on two different / non-overlapping databases to delineate the contours. Note that the Complementarity Plot is probabilistic (knowledge-based) in nature. Based on this strong coherence between the two methods consistent across all three burial bins (CP1, CP2, CP3), the original contours were retained in both (single-dielectric, multi-dielectric) versions of the softwares.

**Fig.2.**
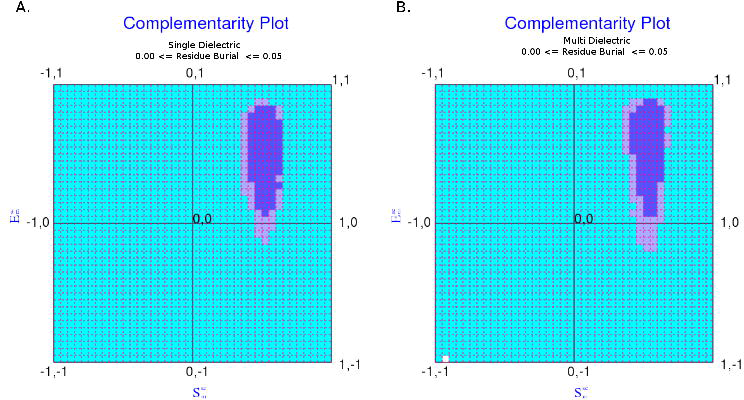
Contours to delineate the different regions (probable, less probable, improbable) in the Complementarity Plot: drawn for CP1 where E_m_ calculated by (A) single‐ and (B) multi-dielectric methods. The ‘probable', ‘less probable’ and ‘improbable’ regions of the plots are colored in purple, mauve and sky blue respectively.

### CP_dock_

In addition to the adaptation of the multi-dielectric method to compute electrostatic complementarity, here we also present an equivalent though new plot (namely, CP_dock_) designed with the view to score docked inter-protein or peptide-protein complexes. CP_dock_ plots single ordered-pair complementarity values computed at the protein-protein interface (Sc, EC) in the background of equivalent non-overlapping contoured regions (i.e., ‘probable', ‘less probable', ‘improbable') delineated from the distribution of the same set of complementarity measures (i.e., Sc, EC) from a database of high resolution protein-protein complexes. In effect, this is a back-track to the origin of the subject since the original idea to evaluate shape and electrostatic complementarities (Sc, EC) in proteins was emerged and implemented for interfaces only, based on single complementarity scores [13,14] assigned to each interface. Here we take the opportunity to demonstrate the numbers pictorially.

To that end, 1880 high resolution protein co-complex crystal structures were assembled from a previously used database (**DB3**, see **Methods**); the (Sc, EC) values calculated at their interface (precisely according to an earlier report [16]) and plotted in accordance with the previous constructions of the plots [1]. The multi-dielectric Gaussian model [12] was implemented to compute EC. The whole two-dimensional area was divided into square grids (0.05 × 0.05 wide) and the probability of finding any point (representative of a protein-protein interface) in a particular grid (P_grid_) was estimated as demonstrated earlier [1]. Likewise to its original constructions [1], the plots were then contoured based on their grid-probability values, P_grid_ >=0.005 for the first contour level and >0.002 for the second. The cumulative probabilities of locating a point within the first and the second contoured level were found to be in strong agreement with previous versions of the complementarity plots constructed to score proteins in a residue wise manner.

One noticeable point of difference between the residue-wise-multi-valued and single-valued versions of the plots is their range in the shape complementarity (**Fig.3**) defined along the horizontal axis. It is to be noted that Sc maps to an elevated range of values in comparison to the residue-wise equivalent measure S_m_ as evident from the two plots. This is because Sc is computed traditionally on the molecular Connolly surfaces [13] whereas S_m_ is computed on van der Wal’s surfaces [18]. It is well-known [18] that the shape complementarity measure renders a higher value when computed on molecular Connolly surfaces than equivalent van der Waal’s surfaces because of the re-entrant points of the van der Waal’s surfaces making their contours (protrusions and crevices) sharper than the corresponding Connolly surfaces [18].

**Fig.3.**
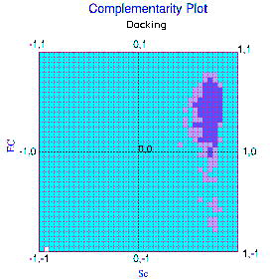
Contours to delineate the different regions (probable, less probable, improbable) in CP_dock_. Sc, EC calculated for interfaces from the database DB3. The ‘probable', ‘less probable’ and ‘improbable’ regions of the plots are colored in purple, mauve and sky blue respectively.

The fact that there is a 20% probability associated with attaining a negative EC at native inter-protein interfaces [16] is also reflected in the isolated islands at the negative half of EC in the plot. The constraints with respect to Sc is again much stronger than EC featured by the horizontal (Sc) and vertical (EC) widths of the corresponding contoured levels (say, the probable regions) in the plot. This difference in relative width (constraints) given rise by Sc and EC is consistent with the short and long range nature of the forces from which they have been originated.

### Utility of CP_dock_ exemplified in a docking benchmark

The apt of the plot (CP_dock_) in discriminating between good and bad realistic protein-protein docked complexes was tested on a recent docking benchmark [20] built using SwarmDock [21] to generate the models. The benchmark (named after the first author as ‘Moal') originally contained 56015 models for 118 targets and was previously used to test, validate and compare the performance of another recent docking scoring function, ProQDock [16] trained by support vector regression machines using shape and electrostatic complementarities as two of its thirteen features. In the same study, it was found that the electrostatic complementarity can as well be negative for top ranked realistic docked protein complexes at a statistically non-negligible fraction (~21%), also true even for native experimentally solved protein complexes (~20%). In such cases, highly raised values of shape complementarity was found to compensate the deleterious electrostatic effect at the interfaces. When Sc, EC were calculated on the native Moal targets, the same fraction (i.e., negative EC) was found to be ~17%, similar to the larger datasets.

Our objective for the current calculation was clear that we wanted to test the efficacy of CP_dock_ in its discriminatory ability between good and bad docking models, only based on the two complementarity measures. If found to be efficient, one can then potentially use this tool as a means to provide a meaningful delineation between good and bad docked models, to be used at the initial screening phase of a docking scoring pipeline, before actually going into expensive computation with more sophisticated trained machines like ProQDock [16]. To that end, one had to consider one or more subsets of (Moal) targets (and their corresponding models) for which their native Sc, EC values were found to be optimal falling in the ‘probable’ / ‘less probable’ regions of the plot. In other words, it probably won’t make a sense to try out CP_dock_ on those targets where the native complex itself maps to the ‘improbable’ region of the plot, due to sub-optimal values being raised by one of its two components (EC in particular, hitting a negative value in ~1/5th of the cases).

Two separate calculations were conducted on two subsets based on two sets of cutoffs. The cutoffs were set according to the normalized frequency (or probability) distributions of the parameters as revealed from the database, **DB3**. The probability distribution of Sc was found to best fit to a sharp Gaussian curve (R2=0.98) with tight bounds on either end of the peak value, while, the same for EC was negatively skewed fitting best to a Lorentzian distribution (R2=0.97) accompanied by a long tail mapping to the negative EC values (**Fig.4**). The nature of the curves were identical to the characteristic patterns depicted earlier for the conditional probability estimates P(S_m_|{Res, Bur}) and P(E_m_|{Res, Bur}) calculated on their corresponding residue-wise parameters S_m_ and E_m_ for a given burial bin (Bur) and residue identity (Res) [1]. The contrasting features in terms of the stringent and relaxed curve-widths of the two distributions were also consistent with the nature of short‐ and long-range forces determining the Sc and EC respectively.

**Fig.4.**
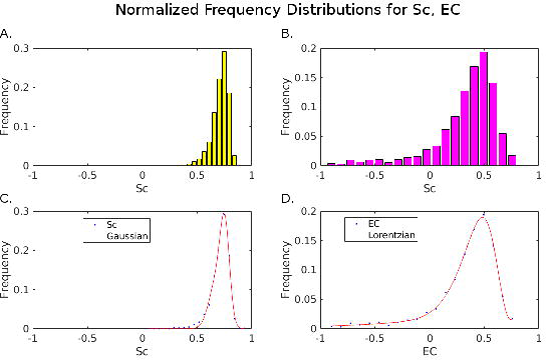
Normalized Frequency (or probability) distributions of (A) Sc and (B) EC and their best fit curves (C) and (D) respectively. The goodness of fit is demonstrated by R2 values of 0.98 and 0.97 for Sc and EC respectively.

To determine the cutoffs based on these two distributions, first, their medians were calculated (µ(Sc): 0.73, µ(EC): 0.41) along with the corresponding absolute median deviations (σ(Sc): 0.06, σ(EC): 0.21). To note is that for EC, the median was certainly a better choice for central tendency because of the negatively skewed nature of the distribution while for Sc, either of the mean or median could have been the choice due to the symmetric nature of the distribution (mean: 0.71, median: 0.73). For consistency purposes, medians were considered for both.

For the first of the two calculations, a straight cutoff (or lower threshold) equaling to the aforementioned median values (Sc: 0.73, EC: 0.41) were set for both measures, while for the second, the corresponding lower thresholds were set to (µ-3σ) and (µ-σ) for Sc and EC, consistent with the stringent and relaxed curve-widths of their respective distributions. Note that the vertical EC-width encompassing the ‘probable’ region is about thrice the horizontal Sc-width in CP_dock_ (also true for the residue-wise CP1 corresponding to the first burial bin).

In effect, only the ‘elite class’ of targets (in terms of Sc, EC) were considered for the first calculation, which could secure highly elevated values of Sc ≥ 0.73, EC ≥ 0.41 in their respective native structures. Only 28 out of 118 targets could fall into this class. While, for the second calculation, those targets were considered for which the native structures could hit values in Sc, EC between closed intervals of the aforementioned lower thresholds (Sc: µ-3σ; EC: µ-σ) and the corresponding medians (µ). 39 targets could fall into this class satisfying the Sc, EC criteria of 0.55 ≤ Sc < 0.73 and 0.20 ≤ EC < 0.41 simultaneously. Hence, this class of targets could be viewed as the ‘average class’ and there’s no overlap between the two ('elite’ and ‘average') classes totaling 67 targets (and the corresponding 31508 models) covered between them.

In effect, for both the ‘elite’ and the ‘average’ class of targets, the native Sc, EC values were optimal, falling into the ‘probable’ or ‘less probable’ regions of CP_dock_, such that they could serve as valid reference native benchmarks (in terms of Sc, EC) against which the quality of the docked models could be judged. We were to test whether in such cases CP_dock_ was indeed able to suggest reasonably strong probability estimates that could delineate between incorrect and correct docked models as assigned to each model by the docking benchmark [20] based on the CAPRI classification scheme [22], in which, correct models refer to the ‘acceptable or higher’ category.

For all targets covered by both the classes (as detailed above) and their corresponding models, Sc, EC were calculated, plotted and overlaid on CP_dock_ with different colors assigned to the native (white), correct (black) and incorrect (red) models. For each target, the fraction of models falling into the ‘probable', ‘less probable’ and ‘improbable’ regions of the plot were calculated separately for the correct and incorrect models. A descriptive statistics on these fractional counts were performed afterwards on the targets falling into the ‘elite’ and ‘average’ class (as defined above) individually.

For the incorrect models belonging to the ‘elite’ class of targets, the fractions (averaged over all targets in the class) falling into the ‘probable’, ‘less probable’ and ‘improbable’ regions of CP_dock_ were 23.0% (±7.4), 37.5% (±3.6) and 39.6% (±7.6) respectively (**Table 1**). In distinct contrast, the same fractions for the correct models in this class were 81.8% (±13.1), 14.9% (±10.8) and 3.3% (±3.7) respectively (**Table 2**).

**Table 1.** CP_dock_ statistics on Incorrect models in the ‘Elite class’ of targets selected from the Docking Benchmark ‘Moal’. F_probable_, F_less_probable_, F_improbable_ stands for the fraction of points falling into the ‘probable’, ‘less probable’ and ‘improbable’ regions of the plots. TotNM stands for the total number of models in the said category (incorrect / correct) as mentioned in the table title.

| **Target** | **F<sub>Probable</sub>** | **F<sub>Less_Probable</sub>** | **F<sub>Improbable</sub>** | **TotNM** |
| --- | --- | --- | --- | --- |
| 1AVX | 0.165 | 0.397 | 0.438 | 424 |
| 1AY7 | 0.134 | 0.459 | 0.408 | 435 |
| 1AZS | 0.186 | 0.355 | 0.459 | 484 |
| 1D6R | 0.232 | 0.424 | 0.344 | 423 |
| 1E6E | 0.248 | 0.368 | 0.385 | 424 |
| 1E96 | 0.229 | 0.361 | 0.410 | 501 |
| 1EAW | 0.470 | 0.394 | 0.136 | 427 |
| 1FCC | 0.257 | 0.371 | 0.371 | 427 |
| 1FQJ | 0.208 | 0.389 | 0.403 | 463 |
| 1GPW | 0.320 | 0.315 | 0.364 | 408 |
| 1GXD | 0.143 | 0.362 | 0.494 | 463 |
| 1I2M | 0.243 | 0.337 | 0.420 | 400 |
| 1J2J | 0.188 | 0.363 | 0.449 | 409 |
| 1JTG | 0.227 | 0.381 | 0.391 | 500 |
| 1K74 | 0.138 | 0.430 | 0.432 | 376 |
| 1KXP | 0.158 | 0.371 | 0.471 | 497 |
| 1LFD | 0.308 | 0.346 | 0.346 | 468 |
| 1ML0 | 0.306 | 0.338 | 0.356 | 488 |
| 1OC0 | 0.123 | 0.380 | 0.498 | 423 |
| 1SYX | 0.286 | 0.423 | 0.291 | 414 |
| 1WDW | 0.158 | 0.303 | 0.539 | 459 |
| 1XQS | 0.262 | 0.379 | 0.359 | 448 |
| 2I25 | 0.292 | 0.333 | 0.375 | 498 |
| 2I9B | 0.268 | 0.409 | 0.323 | 494 |
| 2MTA | 0.245 | 0.378 | 0.378 | 431 |
| 2OZA | 0.181 | 0.409 | 0.409 | 446 |
| 2PCC | 0.214 | 0.331 | 0.455 | 501 |
| 7CEI | 0.241 | 0.382 | 0.377 | 413 |
| Average | 0.230<br>($\pm 0.074^1$) | 0.375<br>($\pm 0.036$) | 0.396<br>($\pm 0.076$) | |

1 Standard Deviation

1 Standard Devision

**Table 2.**
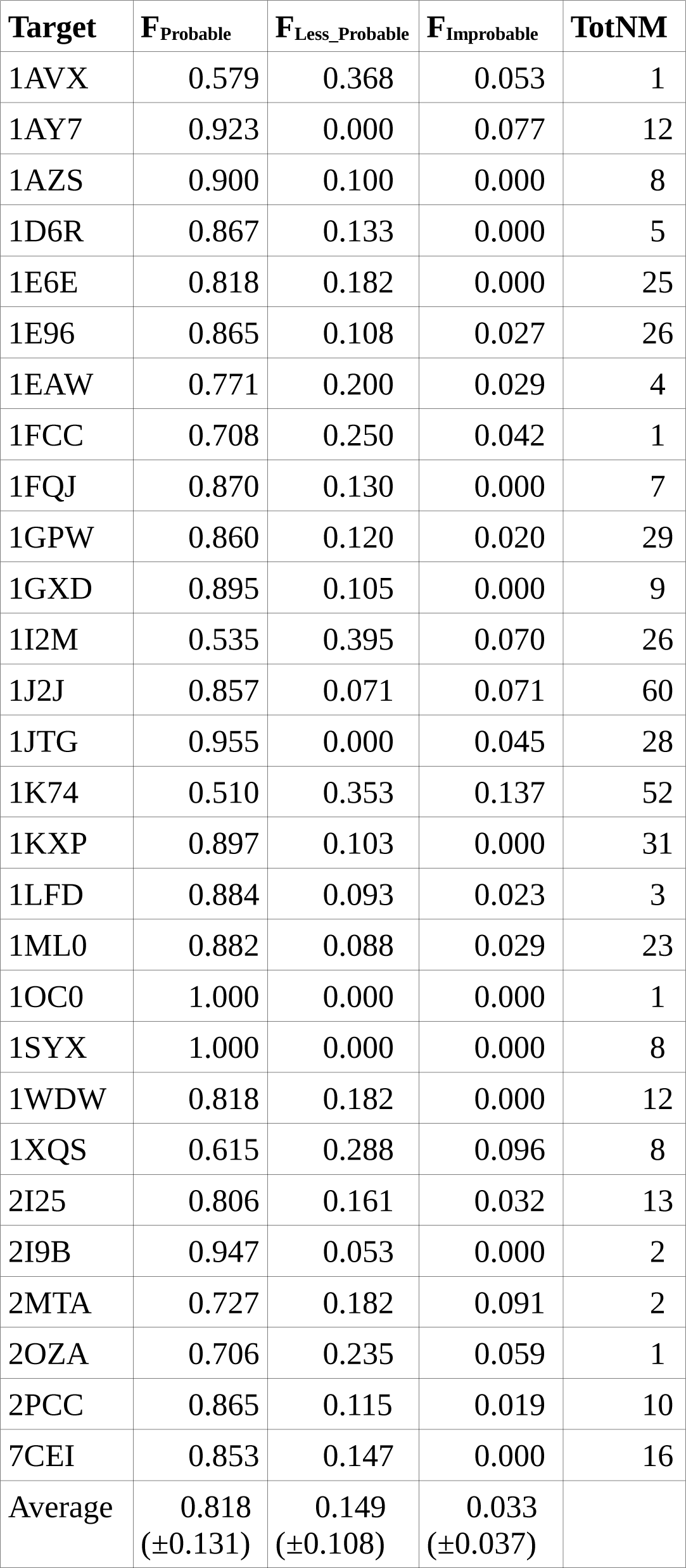
CP_dock_ statistics on correct models in the ‘Elite class’ of targets selected from the Docking Benchmark, Moal. F_probable_, F_less_probable_, F_improbable_ stands for the fraction of points falling into the ‘probable’, ‘less probable’ and ‘improbable’ regions of the plots. TotNM stands for the total number of models in the said category (incorrect / correct) as mentioned in the table title.

A thorough visual examination of the individual plots revealed a few kinds of distribution patterns of the correct models among themselves as well as with respect to their corresponding native(s). In the most prevalent pattern observed, the correct models were found to be clustered together around the closely spaced native (**Fig.5**) mostly falling into the probable regions of the plots. While, another frequent pattern portrayed the distant residence of the native with respect to the cluster formed by the correct models (**Fig.6**). Yet another third pattern was observed where the correct models were themselves scattered, though again mostly falling into the probable regions of the plots (**Fig.7**). On the other hand, the incorrect models were mostly found to be gathering around the left bottom half of the less probable / improbable regions of the plots mapping to lower / sub-optimal Sc, EC values. Overall, the ability of the plot to delineate between the correct and incorrect models were clear and unambiguous in this ‘elite’ class of targets, the ones with optimal and elevated native Sc, EC values providing strong reference points for quality assessment of the docked models.

**Fig.5.**
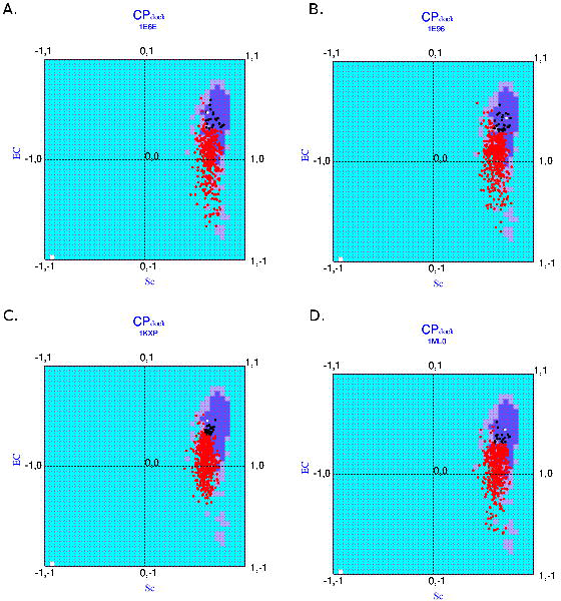
Distribution of points corresponding to the incorrect (red), correct (black) and native structures (white) for the Elite class of targets in CP_dock_: (A) 1E6E, (B) 1E96, (C) 1KXP and (D) 1ML0. These distributions represent cases where the correct models cluster around their corresponding closely spaced natives.

**Fig.6.**
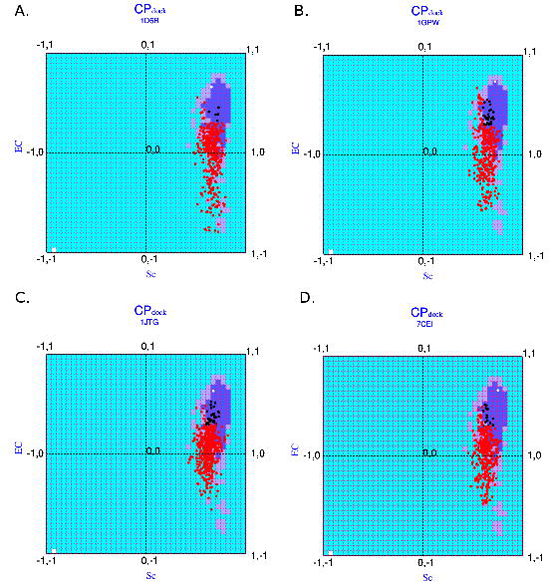
Distribution of points corresponding to the incorrect (red), correct (black) and native structures (white) for the Elite class of targets in CP_dock_: (A) 1D6R, (B) 1GPW, (C) 1JTG and (D) 7CEI. These represent cases where the native is far away from the cluster of the correct models.

**Fig.7.**
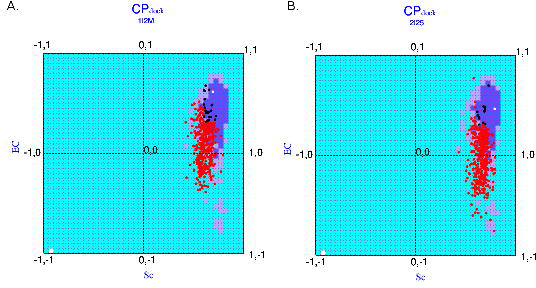
Distribution of points corresponding to the incorrect (red), correct (black) and native structures (white) for the Elite class of targets in CP_dock_: (A) 1L2M, (B) 2I25. These represent cases where the correct models are scattered in contrast to clustering as in Fig.5 and Fig.6.

For the ‘average class’ of targets, the mean fractions of incorrect models falling into the ‘probable’, ‘less probable’ and ‘improbable’ regions of CP_dock_ were 19.6% (±7.2), 35.9% (±3.9) and 44.5% (±9.9) respectively (**Supplementary Table S1**) while the same fractions for the correct models in this class were 64.9% (±12.1), 32.8% (±11.4) and 2.3% (±2.5) respectively (**Supplementary Table S2**).

So to speak, the mean fractional counts (falling into the three disjoint regions of the plots) in both classes (‘elite’ and ‘average’) were fairly similar for incorrect models while for the correct models the ‘elite class’ could hit to a higher mean fractional count in the ‘probable’ regions (81.8%) than to that of the average class (64.9%) with standard deviations of the same order of magnitudes. This has been accompanied by a corresponding lower mean fractional count in the ‘less probable’ regions for the elite class (14.9%) compared to the average class (32.8%) again with roughly identical standard deviations while similar values obtained for both class in the ‘improbable regions’ of the plot. This third category of points could be treated as ‘false negatives’. The relative alterations in the corresponding mean fractional counts in the two class could as well be viewed as the transition of ~15% of points from the ‘probable’ to the ‘less probable’ region, brought about by switching from the ‘elite’ to the ‘average’ class. Note that the native Sc, EC values for the targets in the ‘average class’ is lower in magnitude to that of the ‘elite class’.

Although, the raw numbers suggest that the discriminatory ability of CP_dock_ in the ‘elite class’ is somewhat better than the ‘average class’, if viewed against their corresponding ranges of native reference values (Sc, EC), their performances might as well be interpreted as equivalent in both classes. Also, the dependence of the discriminatory ability of the methodology on the native (Sc, EC) reference values was evident. This said, CP_dock_ indeed appears to be more than useful to be incorporated in the initial screening phase of a docking scoring pipeline, especially effective for those targets which procure strong native reference values in their Sc, EC.

## Conclusions

This brief software report essentially presents an upgraded version of the Complementarity Plot (CP), an established structure validation tool for proteins, based on the incorporation of the multi-dielectric functionality (as in the advanced Delphi version: 6.2 or higher) to make it physico-chemically more realistic. It also highlights the agreement between the two methods (single‐ and multi-dielectric) in terms of both residue-wise (E_m_) as well as single-valued (EC) terms of electrostatic complementarity measures (computed both at protein interiors and interfaces). Hence, it was felt wise to keep both provisions in the software suite. In effect, the conclusions also support the previous observations that the impact of a variety of values chosen as the internal dielectric (in continuum electrostatic models) is rather passive on the electrostatic complementarity, a key determinant of the global long-range harmony in proteins. Having said that, we definitely recommend our users to switch to the more advanced multi-dielectric version of CP for having stronger theoretical foundations potentially giving rise to more reliable E_m_ / EC values. The difference between the E_m_ / EC values computed by the two methods might actually be meaningful in certain structural contexts, and, in such cases some local salient features might be portrayed by the advanced method that might otherwise be missed by the older method (e.g., in mutational studies, protein / peptide design). A full-scale analysis of this kind might serve the subject matter of future studies.

The current report also presents a new methodology and a variant plot, namely, CP_dock_, based on the same principles of complementarity, specifically designed to be used in protein-protein docking. Calculations on a recent *state-of-the-art* docking benchmark reveals the fact that CP_dock_ can indeed be effectively used in the initial screening phase of a docking scoring pipeline, particularly effective for targets with strong native reference values in the two complementarity measures.

## Acknowledgement and Funding

We take the opportunity to acknowledge Prof. Parbati Biswas (Department of Chemistry, University of Delhi) for her support during the course of the current study. The work was supported by the Department of Science and Technology – Science and Engineering Research Board (DST-SERB research grant **PDF/2015/001079/LS).**

